# Hebbian learning accounts for the effects of experience and novel exposure on representational drift in CA1 of hippocampus

**DOI:** 10.1101/2025.10.21.683686

**Authors:** Gloria Cecchini, Alex Roxin

**Affiliations:** Centre de Recerca Matemàtica

**Author notes:** **For correspondence:** (AR).

## Abstract

Neuronal activity patterns slowly change over time even when sensory stimuli and animal behavior remain stable, a phenomenon known as representational drift. In area CA1 of the hippocampus, the amount of drift in the tuning of place cells on a familiar linear track is proportional to the time the animal spends exploring that track while the drift in cells’ mean rate depends on the absolute time elapsed between sessions. Recently it was shown that exploration of a novel, enriched environment between sessions on the familiar track actually decreases the drift in tuning on that track, i.e. it stabilizes place fields compared to a baseline condition without the novel learning. This finding challenges computational models of drift in which new learning leads to overwriting and hence one would expect more drift, not less. Here we show that such models are indeed compatible with the observed findings as long as spatially-tuned input populations to CA1 which are active on the track versus the enriched environment are largely non-overlapping. Furthermore, in order to reproduce the findings, we must assume that the total amount of learning between the baseline and novel-exposure conditions is the same. Namely, the synaptic resources available for encoding patterns, or episodes, is fixed over a given period of time, but can be preferentially allocated to episodes of particular salience, such as the exploration of novel environments.

## Introduction

Chronic, long-term recordings in awake behaving mice have revealed that patterns of neuronal activity, seemingly stable on the timescale of seconds to minutes, nonetheless slowly evolve over the span of hours, days and weeks, even when the associated behavior is stable. This phenomenon, dubbed representational drift (RD), has been observed in a number of cortical areas, including hippocampus ***Mankin et al. (2012***); ***Ziv et al. (2013***); ***Mankin et al. (2015***); ***Rubin et al. (2015***); ***Sheintuch et al. (2017***); ***Rubin et al. (2019***); ***Gonzalez et al. (2019***); ***Levy et al. (2020***); ***Keinath et al. (2021***); ***Khatib et al. (2012***); ***Sheintuch et al. (2023***); ***de Snoo et al. (2023***); ***Geva et al. (2023***); ***Elyasaf et al. (2024***); ***Climer et al. (2025***); ***McLachlan et al. (2025***); ***Madar et al. (2025***); ***Kanter et al. (2025***), parietal cortex ***Driscoll et al. (2017***), motor cortex ***Gallego et al. (2020***), visual cortex ***Deitch et al. (2021***); ***Marks and Goard (2021***); ***Bauer et al. (2024***); ***Brown and McGee (2025***); ***Sotomayor-Gómez et al. (2025***), piriform cortex ***Schoonover et al. (2021***), barrel cortex ***Ahmed et al. (2024***) and auditory cortex ***Noda et al. (2025***).

In a previous study we characterized the role of synaptic volatility and plasticity on RD, primarily in hippocampus, in a computational network model with feedforward architecture ***Devalle et al. (2025***). We showed that random rewiring of synaptic inputs to CA1 pyramidal cells from CA3 and enthorinal cortex (EC) accounted for experimentally observed statistics on RD, including the bi-phasic decrease in the correlation of the population vector, the gradual diffusion of place field centroids and the heterogeneity in the stability of cells over sessions. The session-to-session changes in the synaptic inputs required to fit the data, largely quantitatively, followed simple Gaussian statistics. These statistics were compatible with both undirected synaptic volatility as well as a Hebbian plasticity mechanism for the encoding of random patterns of activity. Thus at the level of the stationary RD observed in CA1 pyramidal cells we could not distinguish between these two synaptic mechanisms.

However, additional experimental findings on the influence of experience on RD pointed to the importance of the Hebbian mechanism. Specifically, when mice were exposed to two distinct environments with differing frequency, one twice as often as the other, it was found that drift in firing rates only depended on the time elapsed between sessions, while drift in place cell tuning depended mainly on time spent on the track ***Geva et al. (2023***). Our computational model with Hebbian plasticity reproduced these results with the assumption that other episodes, unrelated to and uncorrelated with the two explored tracks, were encoded in the network between sessions. Such episodes could be related to the animal’s home cage or other familiar events, much like how human beings store daily episodes even in familiar environments and even within a set routine.

In a subsequent set of experiments, researchers sought to investigate the effect of out-of-session learning on RD explicitly ***Elyasaf et al. (2024***). As a baseline condition, recordings were made for several days without additional learning and only then were animals allowed to explore an enriched environment between sessions on the linear track. Comparison of RD between the baseline condition and that with the novel exposure revealed no effect on drift in mean firing rates. On the other hand, drift in the tuning of place cells in CA1 was actually found to be less with the novel exposure. This was taken as evidence against ongoing learning as the mechanism underlying RD, as one would intuitively assume it ought to have *increased* drift. In this brief manuscript we argue that this finding is, in fact, compatible with ongoing learning, but does provide a strong constraint on how synaptic resources are allocated during the storage of episodes. Specifically, we reproduced the findings from ***Elyasaf et al. (2024***) with the same network model used to reproduce the results from ***Geva et al. (2023***) and, surprisingly, with precisely the same parameter values. The only difference was the correlation in the place-cell inputs between the linear track and the enriched environment, compared to the linear track and other episodes, modeled as random patterns. Importantly, the model was only consistent with experiment if the number of patterns encoded between episodes, and hence the totality of synaptic resources available for learning, was constant in time.

## Results

We modeled the ongoing encoding of patterns in a feedforward network with two input layers. The network model consisted of a layer of linear threshold rate neurons, interpreted as CA1 pyramidal cells, which received input from a layer of spatially tuned cells and a separate layer of untuned cells, which we interpreted as CA3 and EC respectively, Fig.1a-b, see Methods and Materials for a detailed description. For simplicity the size of each layer was *N*. The binary synapses from CA3 to CA1 and from EC to CA1 were plastic and were updated discretely in time according to a Hebbian rule, Fig.1c. At each time step we considered three, sparse random binary vectors for the activity patterns to be encoded in CA1, CA3 and EC. Once the network had been trained on a sufficiently large number of patterns, the model reached a steady state in which individual synapses continuously changed state in time but the matrix statistics remained stationary. We then selected one particular pattern and tracked it in time. After encoding this pattern with the Hebbian rule, we encoded *k* other patterns and then repeated the tracked pattern. The random patterns represented episodes unrelated to the experimental session. Because the effect of these random patterns on drift was equivalent to that of random synaptic volatility ***Devalle et al. (2025***) we cannot distinguish between the two mechanisms. The encoding of the tracked pattern was meant to model plasticity during the experimental session and hence a repetition represented a subsequent session.

**Figure 1.**
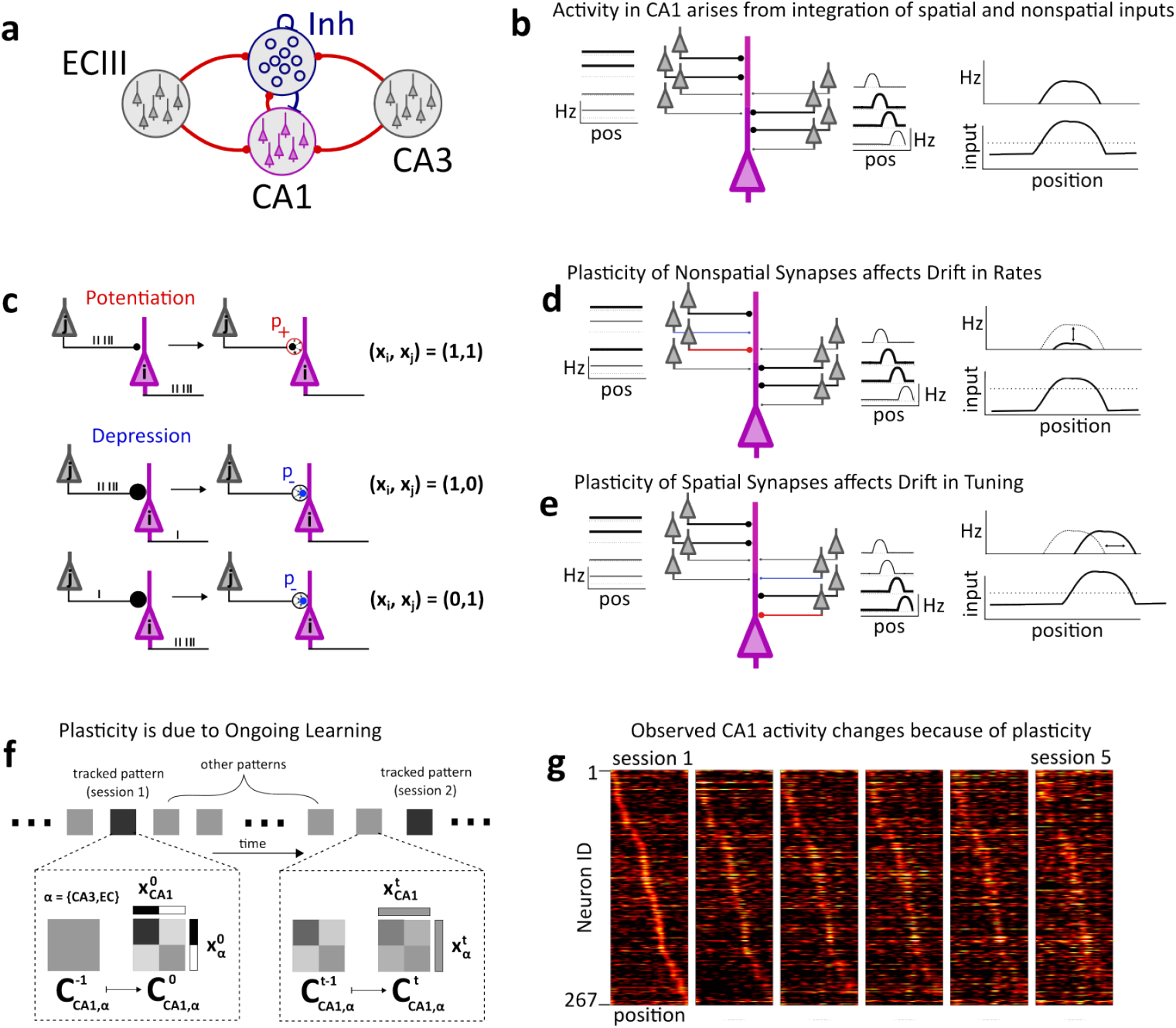
A network model of CA1 accounts for representational drift through ongoing learning. a. Schematic of network architecture. b. Place cell activity in CA1 cells arise through integration of inputs from CA3 and ECIII. The violet cell represents a single CA1 pyramidal cell, which receives inputs via binary synapses, which can be potentiated (depressed) as indicated by thick (thin) lines. ECIII inputs are assumed spatially untuned while CA3 inputs are spatially tuned with uniformly distributed phases. The total input (bottom right) is passed through a threshold linear function to generate the cell output (top right). c. Synapses change according to a Hebbian plasticity rule, where *x*_*i*_ ∈ {0, 1} is the activity of cell *i*. Coactive cells lead to potentiation of the synapses with probability *p*_+_ (top) while depression occurs with probability *p*_−_ when activity levels are different for pre- and post-synaptic cells (bottom). d.-e. The integration of spatially tuned and untuned inputs shown in b. combined with the plasticity rule in c. provide a simple mechanism accounting for changes in the firing rate of CA1 cells (d.) as well as changes in their tuning (e.). f. The fundamental assumption underlying this work is that drift occurs not only through plasticity due to the animal’s experience during the experimental session (black squares), but crucially also through memory storage *between* experimental sessions (grey squares). Specifically, experience during the session leads to the emergence of structure in the connectivity matrix (bottom left) which however can be partially overwritten by the storage of patterns unrelated to the experience (bottom right). g. A sample simulation of a network of CA1 cells with Hebbian plasticity showing the stereotypical drift observed in experiment, sorted by place field location in session 1.

Synaptic plasticity in our model network generated RD because, given identical input patterns at two points in time, changes in the connectivity matrix resulted in distinct output patterns. Specifically, changes in spatially untuned synapses drove drift in firing rates, Fig.1d, while changes in spatially tuned synapses primarily caused drift in cells’ tuning, Fig.1e. Crucially, a key feature of our model is the encoding of the *k* patterns, in general random and uncorrelated with the tracked pattern, between repetitions of the tracked pattern, Fig.1f. The plasticity which occurred while encoding these patterns partially overwrote the structure in the connectivity matrix due to the tracked pattern, leading to drift, Fig.1f (dashed boxes). We interpreted the encoding of these random patterns as the storage of episodic memories between experimental sessions. This resulted in a gradual drift of the population code, reminiscent of the experimentally observed phenomenon of RD ***Ziv et al. (2013***), Fig.1g and ***Devalle et al. (2025***).

### Hebbian plasticity accounts for the effect of experience on RD

In recent work, experiments were conducted in which mice explored two distinct environments, A and B, in an alternating fashion and with differing frequencies ***Geva et al. (2023***). The authors observed that the drift in firing rates in these environments depended on the absolute time between sessions, while the drift in tuning depended on the number of sessions. Therefore, the more experience an animal had on a particular track, the more the tuning of place cells changed. We reproduced this result in our network with Hebbian plasticity by modeling the encoding of patterns of activity corresponding to either track A or B, interleaved with other random and uncorrelated patterns, corresponding to a combination of unrelated episodes, black and grey squares in Fig.2a respectively. In our model, the random overwriting of synapses from spatially untuned inputs was proportional to the number of patterns between sessions, leading the drift in firing rates to be proportional to time, Fig.2b top row. On the other hand, by assuming the fraction of active place-cell inputs for any given episode to be low, interference between sessions was small, and plasticity occurred mainly during a session. Allowing additionally for drift in the place cell inputs, as observed in experiment ***Sheintuch et al. (2023***), further enhanced the contribution of plasticity during the session on drift, compared to other episodes. The drift in the spatially tuned inputs for interleaved patterns did not affect drift as these patterns were already random and uncorrelated with the tracked pattern. The combined mechanisms of significant overlap in spatial inputs only during sessions and additional drift in inputs led to spatial tuning depending on the number of sessions, and not on time, Fig.2b, bottom row.

**Figure 2.**
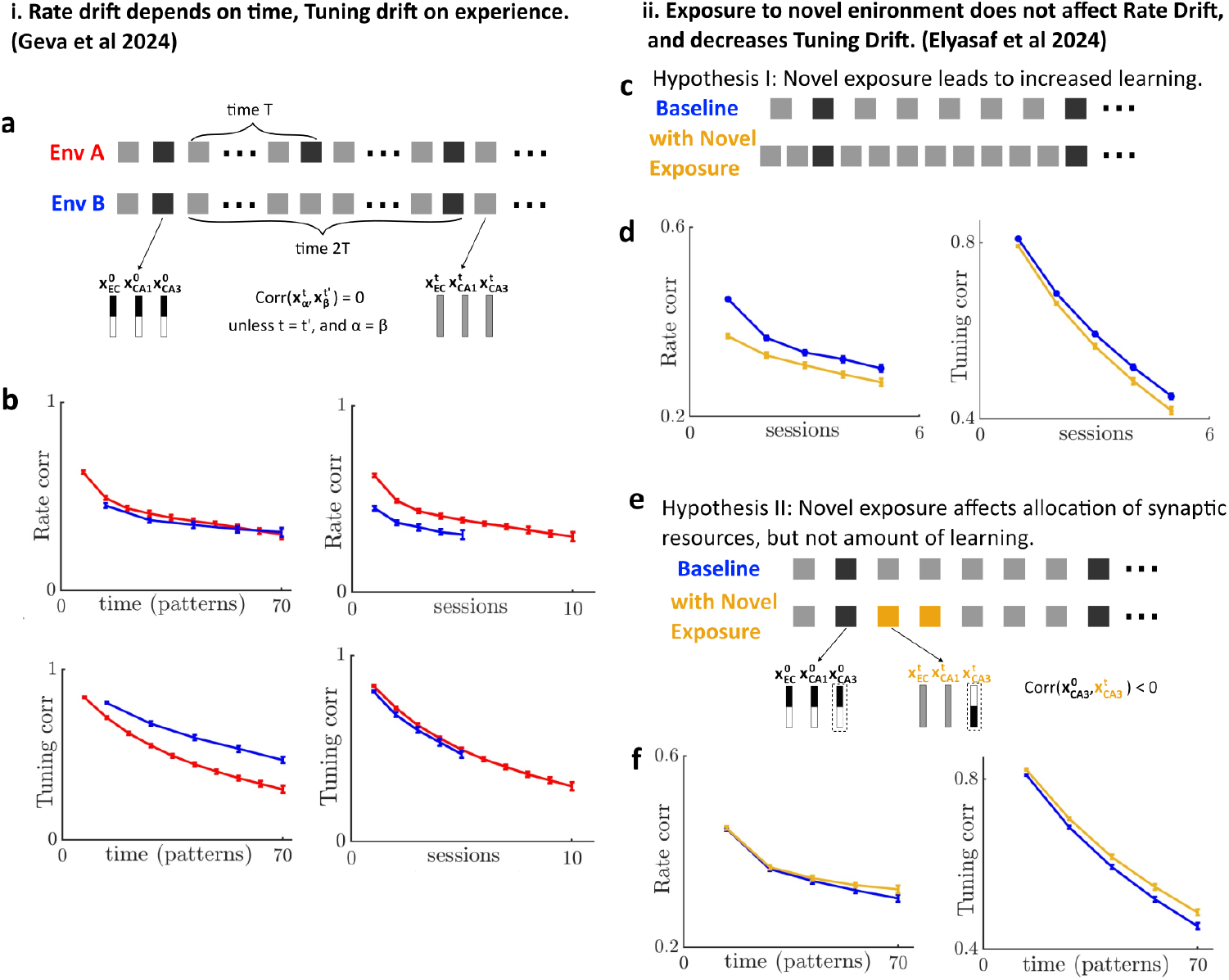
Ongoing learning accounts for the effect of experience on drift in CA1. a. The protocol used in simulation to mimic the experiments from ***Geva et al. (2023***). Environment A is experienced twice as often as environment B, hence the number of random patterns encoded between sessions is greater for environment B. b. In simulations, as in experiment, The drift in firing rates depends on time, here measured in patters (top left) while the drift in tuning depends on the number of sessions (repetitions). c. One possible hypothesis for the effect of novelty in the experiments in ***Elyasaf et al. (2024***). Here we assume that when animals are exposed to a novel, enriched environment between sessions (dark square) the amount of learning increases, i.e. the number of random patterns encoded between repetitions (grey squares) increases. d. Assuming more learning occurs in the face of novel exposure leads to increased drift for firing rates and tuning, in disagreement with experimental findings. e. An alternative hypothesis for reproducing the experimental findings. Here we assume that the amount of learning depends solely on time, and hence the number of patterns stored between sessions is fixed, both for baseline and when there is novel exposure. Rather, the difference between conditions is how the synaptic resources available for plasticity are allocated: randomly for baseline, while with novel exposure place cell inputs for the tracked pattern (linear track) and novel environment (interleaved patterns) are non-overlapping. f. When the total amount of learning is fixed, there is no significant difference between drift in the firing rates (left). Since plasticity in non-overlapping place cell inputs generates less interference than random overlap there is actually less drift in the face of novel exposure (right).

### Hebbian plasticity with fixed synaptic resources accounts for the effect of novel exposure on RD

In order to directly test the effect of out-of-session learning on RD, mice were allowed to explore an enriched arena between sessions on a linear track ***Elyasaf et al. (2024***). In contrast with the expectation that more learning ought to lead to more RD, it was found that exploration of the enriched environment led to *less* drift in place cell tuning compared to baseline, and that the mean rates were unaffected. We sought to model this effect in our network model. For simplicity we used precisely the same parameter values as in Fig.2a-b. We first made the intuitive and naive assumption that exploration of the enriched environment would induce more learning compared to the baseline condition. In our model this is equivalent to increasing the number of random, uncorrelated patterns in between repeats of the tracked pattern, Fig2c. This led to increased drift in both mean rates and cell tuning, as expected, and in contrast with experimental findings, Fig.2d. Next, and guided by the finding that mean rates were unaffected by the novel exposure of the enriched environment, we assumed that the total amount of learning between sessions was fixed. In our model this is equivalent to fixing the number of patterns encoded between repetitions of the tracked pattern, Fig.2e. Furthermore, we hypothesized that the drift in place cell tuning could be less in the novel exposure condition compared to baseline if spatially tuned inputs to CA1 place cells were largely non-overlapping between the linear track and the enriched environment. In biological terms, this would mean that the populations of active CA3 place cells for the linear track and the enriched environment overlap less than that of the linear track with other, familiar environments. In our model this is equivalent to introducing an anti-correlation between the CA3 input vectors for the tracked pattern and some of the interleaved patterns, see Fig.2e. Specifically, for the novel-exposure condition we took 9 of the 13 random input vectors for active CA3 cells to be completely non-overlapping with the tracked input vector. The remaining 4 input vectors were random and uncorrelated as before. Doing this resulted in no significant difference in drift in the rates between the two conditions, while drift in tuning was reduced in the novel-exposure condition compared to baseline in agreement with experiment, Fig.2f.

## Discussion

In this brief manuscript we have shown that the ongoing encoding of patterns in a network model with Hebbian plasticity can account for two highly non-trivial experimental findings regarding the role of experience on representational drift in CA1 ***Geva et al. (2023***); ***Elyasaf et al. (2024***). A key assumption of the model is that plasticity occurs not only due to exploration of the environment during the experimental session, but also between sessions. This same mechanism can account for both the statistics of RD in CA1 pyramidal cells, as well as the effect of repeated exposure to odors on RD in piriform cortex ***Devalle et al. (2025***). In general, the patterns encoded in the model to capture the drift between sessions are chosen to be random and uncorrelated. When this is the case, the storage of these patterns is equivalent to random synaptic volatility (up to second order in the connectivity matrix) ***Devalle et al. (2025***), so both of these mechanisms could underlie some aspects of RD in our model. Indeed, a combination of Hebbian plasticity and synaptic volatility has been shown to account for the statistics of pairwise correlations in auditory cortex ***Eppler et al. (2025***).

Despite the success of models with Hebbian plasticity and synaptic volatility in accounting for data, we believe that the findings in ***Elyasaf et al. (2024***) serve as a cautionary tale. *General* mechanisms which *broadly* agree with experimental data will not be enough to shed light on what causes RD in specific brain areas. Rather, model results must be clearly interpreted in the light of experimental observations, and experiments must challenge theoretical hypotheses. It is only through this back-and-forth between theory and experiment that progress will be made. In the context of our own model, reproducing the findings from ***Elyasaf et al. (2024***) directly constrained how learning occurred between sessions. Interpreting our model results in biological terms, we must hypothesize that the totality of synaptic resources available for learning in the mice in ***Elyasaf et al. (2024***) was a fixed quantity in time. Furthermore, most of these resources were allocated to the learning of the novel enriched environment, and in such a way as to minimize interference of the spatial representation with the familiar track. Whether or not these hypotheses reflect the actual mechanisms underlying RD in those experiments remains to be tested.

## Methods and Materials

We simulated a network model for the activity of pyramidal cells in area CA1 of hippocampus, which was the same one used in our previous work ***Devalle et al. (2025***). The model operated in two modes: 1 - *Plasticity*, and 2 - *Observed Network Activity*.

### 1. Plasticity

At every time step *t* we consider three sparse, random vectors (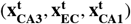) of length (*N*_*CA*3_, *N*_*EC*_, *N*_*CA*1_) which represent the patterns of activity to be stored. For simplicity we take *N*_*CA*3_ = *N*_*CA*1_ = *N*_*EC*_ = *N*. We allow for three possible states of the neurons for plasticity. First, we allow for the possibility that some neurons may not be involved in the plasticity process at all for a given pattern. Therefore for each of the three vectors only a fraction (*f*_*a,CA*3_, *f*_*a,EC*_, *f*_*a,CA*1_) of the cells, chosen randomly, is considered “active”. Synapses with inactive cells are not subject to plasticity. Secondly, of those cells which are active, a fraction *f*_*CA*3_, *f*_*EC*_, *f*_*CA*1_ are considered “high” activity cells, i.e. *x*_*i*_ = 1, while the remaining are “low” activity cells, i.e. *x*_*i*_ = 0. Then, a synapse from cell *j* in population Λ ∈ {*CA*3, *EC*} to cell *i* in *CA*1

1. Is potentiated, i.e. 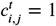, with probability 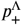 if 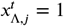 and 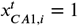. Otherwise 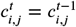.
2. Is depressed, i.e. 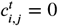, with probability 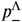 if 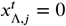 and 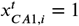. Otherwise 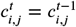.
3. Is depressed, i.e. 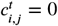, with probability 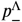 if 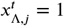 and 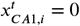. Otherwise 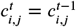.
4. Is unchanged if 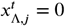 and 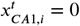, namely 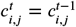.

We repeated this process until the mean and variance of the network connectivity reached a steady state and then tracked a specific random pattern which was presented at a time we defined as *t* = 0, namely (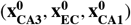). After this time we encoded *k* random, uncorrelated patterns in the network using the plasticity rule above and then repeated a presentation of the tracked pattern. We simulated a series of repetitions of the tracked pattern, always interleaved with *k* random patterns between them. After each repetition of the tracked pattern we simulated the observed neuronal activity as described below.

### 2. Observed Network Activity

We modeled the population activity of CA1 cells as linear threshold units, i.e. the activity of cell *i* at time *t* was given by

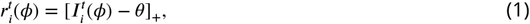

where [*x*]_+_ = *x* if *x >* 0 and is zero otherwise. The input *I*_*i*_ was given by

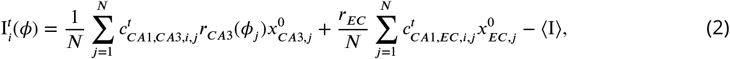

where the rates in CA3 *r*_*CA*3_(*ϕ*) are given by

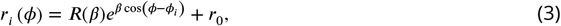

where *ϕ*_*i*_ was the center of the place field of neuron *i, r*_0_ a baseline firing rate, and 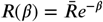 where 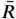 was a constant that set the maximal firing rate. The parameter *β* determined the sharpness of the place fields. The inhibition here was taken into account by subtracting off the mean input, averaged across the network ⟨*I*⟩, and *ϕ* is the position along a circular track. Experimental work has shown that the spatially tuned cells in CA3 themselves undergo some degree of drift (though less than in CA1). To account for this drift we allowed the position of the *i*^th^ place cell (center of the von-Mises distribution) at session *s* to shift with respect to the position at session *s* − 1 according to 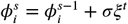, where *ξ* is uniformly distributed from {−*π, π*} and *σ* = 0.025.

We simulated the population activity for each session (repetition of tracked pattern) using Eq.1. We calculated the rate and tuning correlation between the population activities for different sessions as in ***Geva et al. (2023***); ***Devalle et al. (2025***). Briefly, for the rate correlation, we averaged over the spatial variable *ϕ* for each neuron, obtaining a vector of mean rates for each session. We then calculated the correlation of these rate vectors. For the tuning correlation we calculated the correlation of the tuning curve for each cell for a given pair of sessions, obtaining a vector of correlations. We then calculated the mean of this vector.

The parameters were: *f*_*EC*_ = *f*_*CA*1_ = *f*_*CA*3_ = 0.5, *f*_*a,EC*_ = *f*_*a,CA*1_ = 1, *f*_*a,CA*3_ = 0.1, 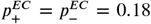, 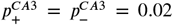, *N* = 1000, *θ* = 0. 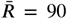, *r*_0_ = 5, *β* = 19, *r*_*EC*_ = 2.25, *σ* = 0.025. For the red curves in Fig.2b, the number of patterns between repetitions of the tracked pattern was *k* = 6 while for all other simulations it was *k* = 13. For all simulations except those in Fig.2f, the patterns between repetitions were random and uncorrelated, between one another and with the tracked patterns. For Fig.2f, the components of the *k* = 13 patterns between repetitions for EC and CA1, namely (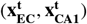), were random and uncorrelated as before. However, the binary activity vector for CA3 was taken to be anti-correlated to that from the repetition for 9 of the 13 patterns. The other 4 were completely random and uncorrelated. The anti-correlation was introduced by taking the fraction of active cells *f*_*a,CA*3_ = 0.1 to be completely non-overlapping with the active cells for the repetition.

## Acknowledgments

This project has received funding from Proyectos De Generación De Conocimiento 2021 (PID2021-124702OB-I00). This work is supported by the Spanish State Research Agency, through the Severo Ochoa and Maria de Maeztu Program for Centers and Units of Excellence in R&D (CEX2020-001084-M). We thank CERCA Programme/Generalitat de Catalunya for institutional support.

